# An inhomogeneous Poisson model of the spatial intensity of opioid overdoses, as represented by EMS naloxone use

**DOI:** 10.1101/529206

**Authors:** Christopher W. Ryan

## Abstract

From 1999 to 2017, the age-adjusted annual death rate in the United States from opioid overdose has increased several-fold, to about 18.3 per 100,000 population. The federal government has declared opioid overdose a public health emergency. Spatially-aware analyses and surveillance may contribute to control of this growing problem.

As naloxone has only one clinical use—the treatment of opioid overdose—its administration by emergency medical services (EMS) personnel can serve as a surveillance indicator for opioid overdose. I previously demonstrated spatial clustering of naloxone-involved EMS calls, over and above that to be expected fromEMS calls in general. To better understand the nature of that clustering, I modelled the spatial distribution of naloxone-involved EMS calls in a three-county region in south-central New York State as an inhomogeneous Poisson process, using as predictors several census-tract-level sociodemographic variables and the point locations of convenience stores (minimarts). In Monte Carlo simulations, I examined how well the model explained the observed clustering.

Although it assumes no interaction between event locations, the inhomogeneous Poisson model was nevertheless able to explain much of the observed spatial clustering. The spatial intensity of EMS calls for opioid overdose, in events per square kilometer, was signficantly higher in census tracts with lower rates of owner-occupancy of housing. Neither the proportion of households in poverty nor the proportion of residents in the 20–44 age band were signficant predictors. Spatial intensity of overdose events decreased by about 10% for each kilometer of distance from the nearest minimart, but this was of borderline signfificance at conventional levels. These findings cast some doubt on the utility of real-time surveillance for apparent spatial clusters of opioid overdose and instead favor a longer-term, systemic strategy comprising efforts to improve neighborhood conditions.

## 3 Introduction

The increasing problem in the United States with opioid dependence and overdose, often fatal, is well-recognized. From 1999 to 2017, the age-adjusted annual death rates from natural and semi-synthetic opioid analgesics, heroin, and non-methadone synthetic opioids (mainly fentanyl and its analogs) have increased by about 400%, 450%, and 3000%, respectively. The increase for heroin has been most marked since about 2010, and that for fentanyl and analogs since about 2013. As of 2017, the age-adjusted death rate from overdose on those opioids stood at about 18.3 per 100,000 population.*^1^*

A variety of indicators can be used in ongoing surveillance of the opioid dependence problem, and of the corollary problem of opioid overdose. Since naloxone is a unique and specific opioid antagonist, administration of naloxone by emergency medical services (EMS) personnel is a potentially useful epidemiological indicator of opioid overdose.

Temporal trends in naloxone use can of course be monitored, but spatial patterns may also be of interest. Published studies of the spatial aspects of pre-hospital naloxone administration have tended to use areal or lattice methods, aggregating counts or rates of events by administrative districts such as states, towns, or census tracts. Examples include work by Merchant and colleagues*^2^* and Klimas and colleagues,*^3^* both of which described a concentration of naloxone-involved EMS calls in a few census tracts, often near the center of large cities.

An areal approach is informative but is subject to the well-known modifiable areal unit problem, in which the observed patterns may vary with the scale to which the data are aggregated. Given that the incident location is usually available in EMS databases, methods for the analysis of spatial point patterns might yield insights unavailable from an areal approach. Using those methods, in a purely descriptive way, I previously demonstrated spatial clustering of naloxone-involved EMS calls, over and above that to be expected from the inevitable clustering of EMS calls to areas of concentrated population.*^4^* The question now turns to why that might be.

Conceptually, an observed pattern of event locations is usually considered to be a realization of a spatial point process that at any given location u has an intensity λ(*u*) in, say, events per *km*^2^. That spatial intensity may be constant across the region (λ(*u*) = λ), but more commonly (and more interesting from an epidemiological standpoint), it may vary from one location to the next, depending on the values of a host of geographic, demographic, or socioeconomic predictors. This describes, broadly speaking, an inhomogenous Poisson model—inhomogeneous because the intensity varies by location, and Poisson because the events occur independently of one another. As a further refinement, a spatial point process may also entail interaction between event locations, such that the occurrence of one increases or decreases the probability of another occuring nearby. These are not Poisson processes. Fundamental analytical tasks with any spatial point process are to model the spatial intensity as a function of scientifically meaningful predictors and to assess the relative contributions of those predictors, and of point-to-point interaction, to the observed spatial heterogeneity.

As an initial modeling step, here I fit an inhomogeneous Poisson model to an observed point pattern of EMS naloxone administration, using as predcitors a number of census-tract-level sociodemographic predictors and the point locations of convenience stores or minimarts. The latter have been associated spatially with other undesirable events*^5^* but have not previously been studied with respect to opioid overdose. While minimarts do not, or course, sell opioids, they are distribution points for a number of other psychoactive substances (e.g. nicotine, caffeine, alcohol) frequently used by people with opioid use disorder, and polysubstance use is common.*^6^* They are also transient waypoints for large numbers of people at all hours of the day and night. I assess model fit, in particular how well the Poisson assumption, not permissive of inter-point interaction, explains the observed clustering.

## 4 Methods

### 4.1 Setting and data sources

The setting for this study is an EMS region comprising three counties in south-central New York State, with a population of about 300,000 and a surface area of about 5500 *km*^2^. The region comprises rural and suburban areas, a number of villages, and two small cities. It is served by 77 EMS agencies of various structures: commercial, volunteer, fire-service-based, transporting, non-transporting, advanced life support (ALS), and basic life support (BLS). Parenteral naloxone has been a standard part of ALS protocols in the region for decades. Recently intranasal naloxone has been widely deployed in the region with BLS personnel, firefighters, and police officers as well, but the study period pre-dates that development.

EMS agencies with a total call volume during the study period of less than 50 were excluded, leaving 63 agencies that were invited to participate by allowing their data to be used.

The Regional Emergency Medical Services Council (REMSCO) maintains an electronic database of patient care reports (ePCR) from all EMS agencies in the Region. The database complies with the National Emergency Medical Services Information System (NEMSIS) standards (http://www.nemsis.org/.) Information about incidents and patients is entered into the database by EMS crews in real time or nearly so. The Regional Emergency Medical Advisory Committee (REMAC—a physician subcommittee of the REMSCO) is charged with stewardship of the database and approved its use for this study. The SUNY Upstate Medical University Institutional Review Board also approved the study.

The REMAC provided a data file consisting of incident locations for all EMS calls handled by participating agencies during the study period of 9 September 2012 to 9 February 2014, inclusive. EMS calls in which naloxone was administered to the patient were considered cases, and other calls were considered controls. Incident locations were geocoded in ArcGIS 10.3, using US Census Bureau TIGERline street address files for the three counties, and county boundary shapefiles from the Cornell University Geospatial Information Repository (CUGIR) at http://cugir.mannlib.cornell.edu/. Match accuracy was set at 79%. No manual re-matching of unmatched or tied addresses was attempted. All geographic objects were projected in Universal Transverse Mercator (UTM) zone 18 N with North American Datum (NAD) 1983 and distances in kilometers. Not unexpectedly, there were a large number of duplicate incident locations among both the cases and the controls. Duplicated locations were removed, meaning that any location to which EMS responded more than once during the study period was represented only once in the data used for analysis.

### 4.2 Model fitting and diagnostics

I fit a Poisson point process “raised incidence” model*^7^* to the spatial intensity of naloxone cases, in events per *km*^2^. This is essentially a log-linear model with an offset, in which predictors have multiplicative effects on the response (Equations 1 and 2).

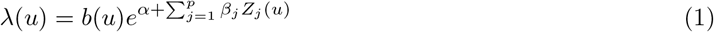

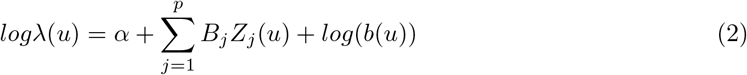

where

- *u* is some spatial location in the region
- there are *j* = 1 …*p* predictors
- *Z_j_*(*u*) represents the value of the *j^th^* predictor at location *u*
- *b*(*u*) is the spatial intensity of the “baseline” events—all EMS calls
- *α* represents the regression intercept

Four predictors were obtained from The US Census Bureau American Fact-Finder website, at the census-tract level, and based on five-year estimates from the 2010-2014 Ameican Community Survey:

- Percent of occupied residences in the census tract that are occupied by their owners
- Percent of households in the census tract in which total household income is below the federal poverty level applicable to the size of the household
- Percent of census tract population that is between the ages of 20 and 44 years (an age range which, informally, encompasses about half the opioid overdoses in the study region)
- Census tract population density in people per *km*^2^

The final predictor was the distance from each naloxone-involved EMS call to the nearest minimart, defined as a co-located gas station and convenience store. Geocoded locations of minimarts were kindly provided as an ESRI shapefile by Dr. Chengbin Deng of the Binghamton University Department of Geography.

Understanding the spatial aspects of any human health event requires a representation of the spatial distribution of the underlying human population—the locations in which the event *could* occur. I calculated a kernel density smoother of the spatial intensity of all EMS calls to represent that underlying density, and used it as an offset in the model, to control for the inhomogenous spatial distribution of the population, and perhaps more to the point, of the population that tends to call ambulances.

**Table 1:** Unmatched, tied, and duplicate locations, and the final analytical dataset.

|  | cases | controls | sum |
| --- | --- | --- | --- |
| locations matched | 198 | 34571 | 34769 |
| locations among multiple matches | 4 | 1449 | 1453 |
| locations unmatched | 45 | 8671 | 8716 |
| unique locations among those matched | 183 | 10643 | 10826 |

In general I followed a purposeful model-building strategy as described by Hosmer, Lemeshow, and Sturdivant.*^8^* I included in the preliminary model all predictors that were statistically significant at the 20% level in univariate testing. Predictors insignificant at the 0.05 level in the multi-predictor model were removed, unless doing so changed the coefficients on remaining predictors by more than 20%, suggesting substantial confounding or collinearity. I assessed the adequacy of the fit of the final model via number of residual-based diagnostics.

### 4.3 Monte Carlo simulations to assess remaining clustering

I then conducted 199 Monte Carlo simulations from the final model, and another 199 from the naive model with no predictors save the offset, to assess how well the final model explained the clustering previously found among the observed naloxone-involved locations. In practice, EMS crews record call locations by street address, and these were the locations used to geocode the observed call location. To mirror this process, the points generated in each simulation were displaced to the nearest road segment. I generated an inhomogeneous K function from each simulated point pattern, and from these samples of K functions, I constructed 95% global acceptance bands under each of those two models. I compared these graphically to the inhomogenous K function from the observed naloxone locations.

## 5 Results

### 5.1 Overview

Thirteen of the 67 eligible EMS agencies agreed to participate. These thirteen comprise the largest and busiest agencies, and together they account for over 90% of total regional call volume.

After eliminating unmatched addresses and reducing duplicated locations to single points, the analytical dataset consisted of 183 case locations and 10,643 control locations (Table 1). Manually matching the case locations that remained unmatched after the automated procedure would have been possible, but this would have been infeasible for the rather large number of control locations, so in an effort to treat cases and controls in the same manner, no manual matching was undertaken. The approximate case and control locations are shown in Figure 1, where the expected heterogeneity is obvious.

### 5.2 Modeling

In fitting a series of univariate Poisson point process models, with the kernel density of all call locations as an offset, each predictor was highly statistically signficant by conventional standards. Therefore all predictors were included in a multivariate “raised incidence model,” with results shown in Table 2.

**Figure 1:**
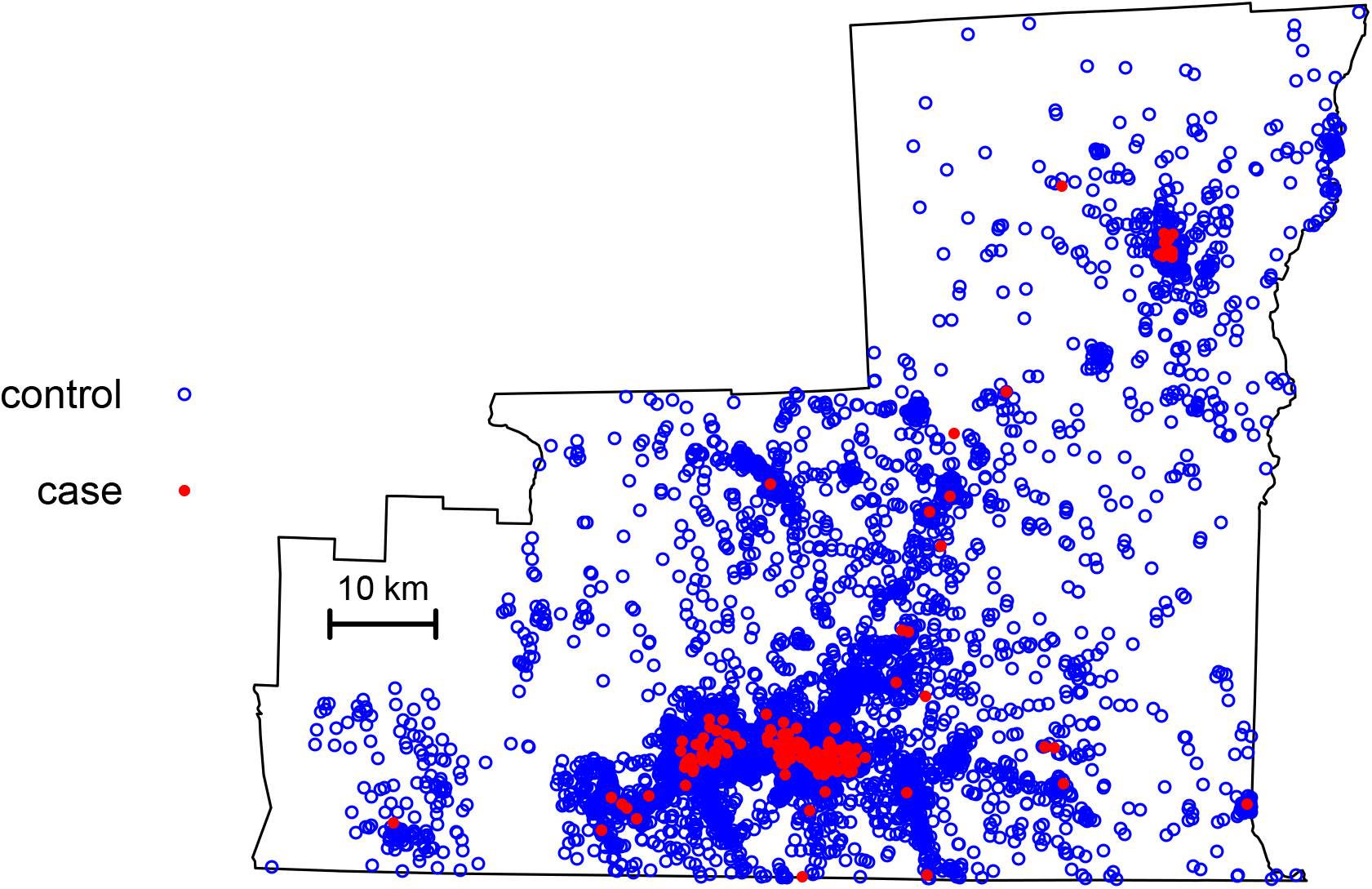
Approximate locations of naloxone cases and non-naloxone controls.

**Table 2:**
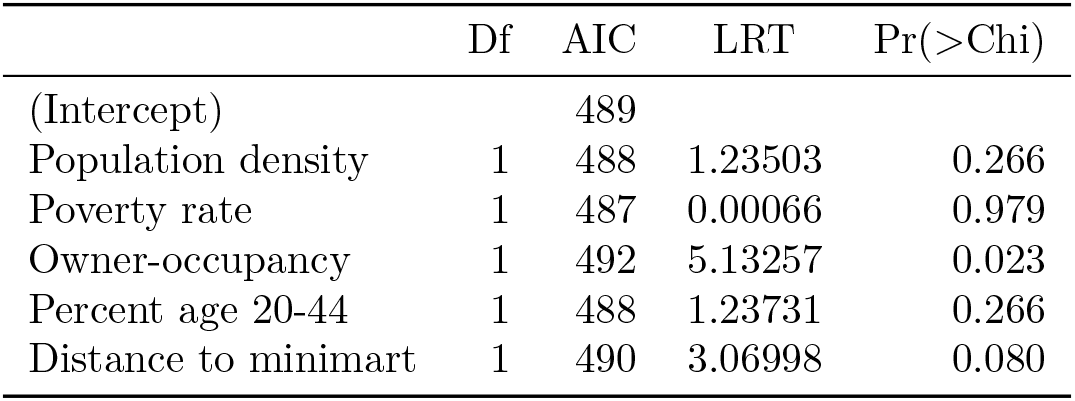
Preliminary multi-predictor model. Df = degrees of freedom for a likelihood ratio test. AIC = Akaki’s Information Criteria if the predictor was removed from the model. LRT = likelihood ratio test statistic. Pr(>Chi) = p-value for the likelihood ratio test.

| | Df | AIC | LRT | $\text{Pr}( > \text{Chi} )$ |
| --- | --- | --- | --- | --- |
| (Intercept) |  | 489 |  |  |
| Population density | 1 | 488 | 1.23503 | 0.266 |
| Poverty rate | 1 | 487 | 0.00066 | 0.979 |
| Owner-occupancy | 1 | 492 | 5.13257 | 0.023 |
| Percent age 20-44 | 1 | 488 | 1.23731 | 0.266 |
| Distance to minimart | 1 | 490 | 3.06998 | 0.080 |

Table 3 shows the percent changes in the remaining coefficients when each predictor alone is removed. As indicated by the very large percentage swings in the coefficients on predictors that remain when one is removed, there was much colinearity, or confounding, between the sociodemographic predictors. This is not unexepected. The only predictor not afflicted by substantial colinearity is poverty rate. Since this predictor was also far from statistically significant in the preliminary multi-predictor model in Table 2, it was removed, resulting in the final model shown in Table 4.

**Table 3:** Percent change in remaining model coefficients upon deletions. Blank cell(s) in each column indicate the predictor that was removed. Column m11 indicates the effect of removing all non-significant predictors from the preliminary model simultaneously. The only removal that leaves the remaining coefficients little-changed is that of poverty rate, shown in column m7

|  | m5 | m6 | m7 | m8 | m9 | m10 | m11 |
| --- | --- | --- | --- | --- | --- | --- | --- |
| (Intercept) | 0 | 9.9 | -0.5 | 52 | -22 | -6.1 | -32 |
| Population density | 0 |  | 1.4 | -40 | 44 | 39.4 |  |
| Poverty rate | 0 | 2611.1 |  | 7083 | -1709 | -1835.7 |  |
| Owner-occupancy | 0 | -11.0 | 0.9 |  | 21 | 28.0 | 59 |
| Percent age 20-44 | 0 | 44.4 | -0.9 | 89 |  | 0.5 |  |
| Distance to minimart | 0 | 16.3 | -0.4 | 52 | 1 |  |  |

**Figure 2:**
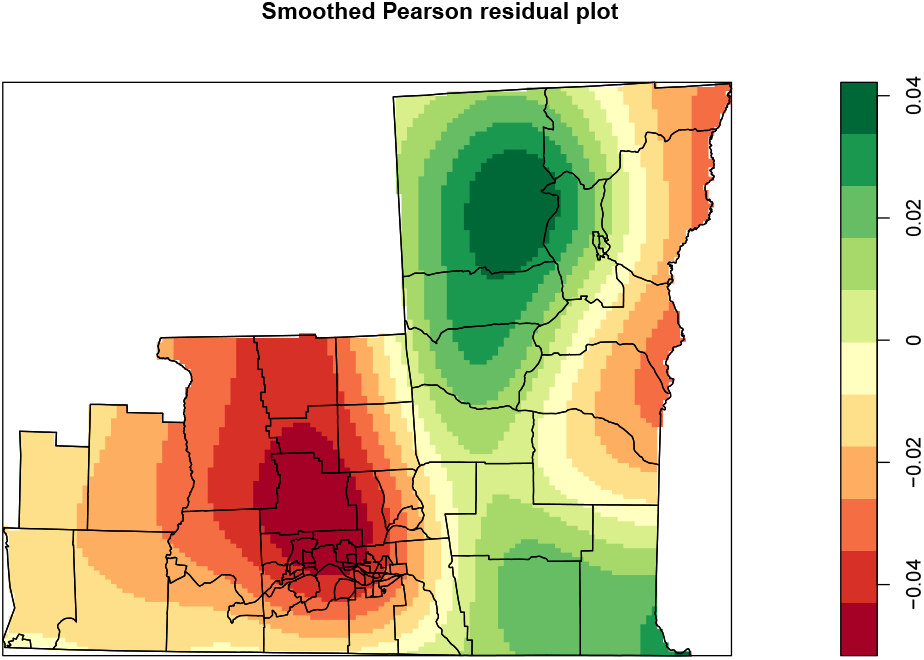
Smoothed Pearson residuals from the final model in Table 4. Mininal under- (green) and over- (red) prediction is seen, but the magnitude of most residuals is less than 0.04 which, at twice their standard deviation, would be considered to indicate a reasonable fit.

**Table 4:** Final model. CI = confidence interval. LB = lower bound. UB = upper bound.

|  | Odds ratio | 95% CI |  |
| --- | --- | --- | --- |
|  |  | LB | UB |
| (Intercept) | 0.028 | 0.009 | 0.088 |
| Population density | 1.000 | 1.000 | 1.000 |
| Owner-occupancy | 0.162 | 0.065 | 0.407 |
| Percent age 20-44 | 5.059 | 0.364 | 70.348 |
| Distance to minimart | 0.897 | 0.788 | 1.020 |

### 5.3 Model diagnostics

A plot of smoothed Pearson residuals from the final model (Table 4) is shown in Figure 2. There are areas of slight over and under prediction, but the magnitude of the residuals is generally less than twice their standard deviation, which would be considered a reasonable fit.*^7^*

### 5.4 Monte Carlo simulations to assess clustering

In Figures 3 and 4, the inhomogeneous K function from the observed case locations is compared against the simulation envelopes from the naive model with the offset but no predictors (left) and those from the final model (right). Deviations of the observed K function from the case locations outside of the 5% critical simulation band suggests clustering that is unexplained by the model used for the simulation. The naive model leaves much of the observed clustering unexplained. The final model, which includes sociodemographic predictors at the census tract level plus the point locations of minimarts, better explains the observed clustering. However, even with the final model, some clustering remains unexplained, as indicated by the observed K function drifting slightly above the simulation band.

**Figure 3:**
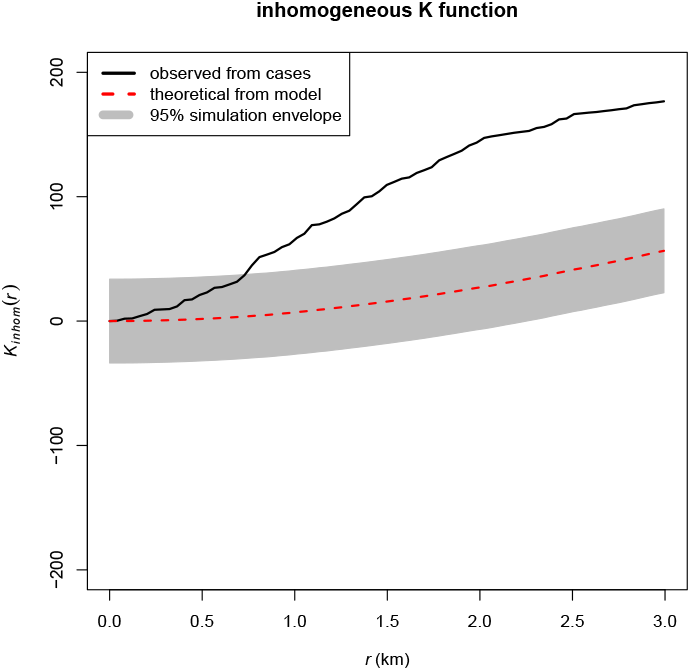
Simulation of inhomogeneous K function from the naive model, with offset only.

**Figure 4:**
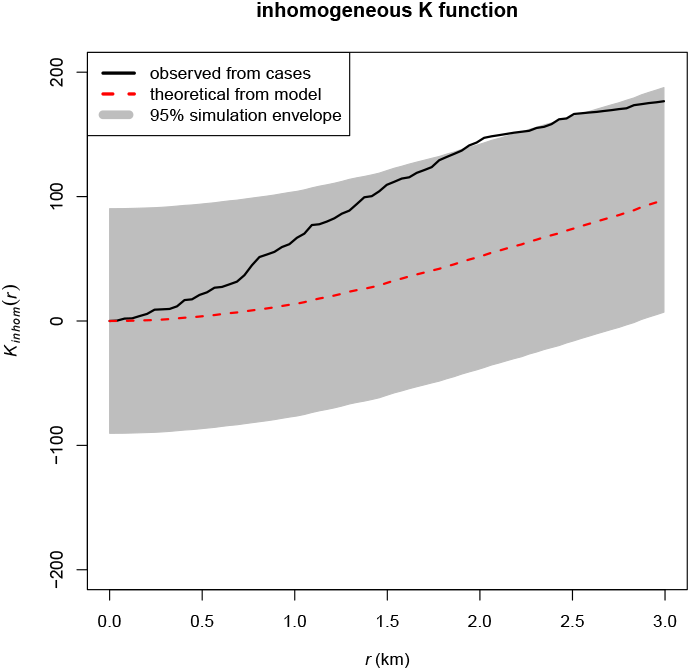
Simulation of inhomogeneous K function from the full multi-predictor model.

## 6 Discussion and limitations

In previous descriptive work, I demonstrated spatial clustering of naloxone-involved EMS calls (a surrogate for opioid overdoses) beyond that to be expected among EMS calls in general.*^4^* The mechanisms underlying observed spatial clustering of a health event carry important implications for public health action. Notionally, these mechanisms span a spectrum from:

1. physical, demographic, or socioeconomic characteristics of neighborhoods that might attract events, with no interaction or attraction between event locations, to
2. attraction between points, such that the occurrence of an event in a particular location stimulates the occurrence of additional events nearby, irrespective of surrounding neighborhood conditions.

There is likely a broad range of hybrid mechanisms between those two “pure” extremes. Public health action on opioid overdose can be guided by the relative contributions of those mechanisms. If the first mechamism predominates, then long-term systemic efforts to improve socioeconomic and physical conditions in neighborhoods would be most important. Resource-intensive efforts to respond “in real time” to apparent spatial clusters of opioid overdose, in the hopes of stopping their expansion, would not be necessary. On the other hand, if the second mechanism predominates, then such real-time intervention activities may be warranted.

The inhomogeneous Poisson model presented here assumes no interaction between event locations but was able to explain much of the observed spatial clustering using neighborhood characteristics. The spatial intensity of EMS calls for opioid overdose, in events per square kilometer, was signficantly higher in census tracts with lower rates of owner-occupancy of housing. Neither the proportion of households in poverty nor the proportion of residents in the 20-44 age band were signficant predictors. Spatial intensity of overdose events decreased by about 10% for each kilometer of distance from the nearest minimart, but this was of borderline signfificance at conventional levels.

This casts some doubt on the need for real-time surveillance for, and response to, apparent spatial clusters of opioid overdose and instead favors a longer-term, systemic strategy to improve neighborhood conditions. The results presented here might nominate measures to increase home ownership, and changes to zoning regulations to reduce the density of minimarts.

There is some precedent for the notion that physical and retail landscape features can influence the spatial intensity of undesired events: Xu and colleagues in Newark, NJ, considering predictor locations and event locations as points on a street network rather than on an unbounded plane, found that shootings tended to cluster near liquor stores, bus stops, grocery stores, and foreclosed houses.*^5^* They found no relation with gas stations. However, it was not clear from their report whether the categories for the retail establishments were mutually exclusive. Thus it is unclear how they would classify minimarts, which in my study area are usually part gas station, part liquor store, and part limited-inventory grocery story.

The distinction between clustering and clusters should be noted. This was an effort to explain global clustering of naloxone-involved EMS calls; it was not a search for specific clusters. As Besag and Newell explain, “In tests of clustering, the usual question is whether the observed pattern of disease in one or more geographical regions could reasonably have arisen by chance alone.” In the present study, the null hypothesis of “chance alone” is adjusted for the spatial distribution of all EMS calls, for census tract sociodemographic variables, and for point locations of minimarts. In contrast, the purpose of cluster detection is to search, perhaps repeatedly, “… a very large expanse of small administrative zones for evidence of individual ‘hot spots’ of disease …”.*^9^* Both strategies are useful, but for different purposes.

Several limitations, both conceptual and technical, must be considered. This was an observational study and thus may not lend itself to causal inferences. Absent the ability to conduct randomized controlled trials of neighborhood characteristics and the built environment, causal evidence of their effects on opioid overdose may be difficult to obtain.

This study constructed and used a non-parametric kernel density estimator of the spatial intensity of all EMS calls. Constructing such an estimator involves art as well as science, mainly mediated through decisions about bandwidth and choice of kernel function. Constructing a consistent kernel density estimator may be particularly challenging in a highly complex urban environment. Recently developed analytical strategies that do not require this step present a new and perhaps better analytical option.*^10^*

The representation of all call locations as street addresses, even if the patient was found far from a street (e.g. under a tractor out in a farm field) presented a challenge. However, projecting the Monte Carlo sampled locations to the nearest road is a reasonable solution and has the advantage of reflecting real-world EMS practice (e.g. recording incident location as the farm’s address). My analysis considered space as an unbounded plane, and this may have some weaknesses, as discussed by Xu, et al,*^5^* particularly since the “urban core” of my diverse study region is trisected by the confluence of two rivers.

Participation by EMS agencies was not universal; however, the vast bulk of EMS calls in the region was included. Duplicate locations were eliminated, mainly to justify the use of relatively simple methods to assess clustering. Thus the analytical data set comprised locations where an ambulance had responded at least once during the study period. Methods that accomodate duplicate locations may yield different results.

My analytical dataset did not contain information on when each event occurred. Thus my analysis is time-agnostic and cannot speak to the question of which events occurred after a nearby event.

Automated geocoding was incomplete, with many locations left unmatched. While manual matching could have been undertaken with the unmatched case locations, it would have been impractical to do the same for the much larger number of unmatched control locations (see Table 1). I believe the methodologic rigor of treating cases and controls in the same manner outweighed any value to be obtained from manually matching the remaining case locations to enable their inclusion in the dataset.

## Supporting information

R code used for anlaysis (with filename extension changed to .txt so as to work with BioRxiv upload)

## 7 Acknowledgements

I thank the following individuals and agencies for valuable assistance with this project:

- Jillian Shotwell, MPH, for geocoding
- Professor Chengbin Deng of the Binghamton University Department of Geography for geocoded locations of minimarts
- The dedicated EMS providers in the Susquehanna EMS Region—skilled clinicians, and the best field epidemiology team one could ask for.

## References

1. Hedegaard, H., Minino, A. M., Warner, M., “Drug Overdose Deaths in the United States, 1999–2017”, NCHS Data Brief 329 (National Center for Heatlth Statistics, 2018), (2018; https://www.cdc.gov/nchs/data/databriefs/db329-h.pdf).

2. Merchant, R. C., Schwartzapfel, B. L., Wolf, F. A., Li, W., Carlson, L., Rich, J. D., Demographic, geographic, and temporal patterns of ambulance runs for suspected opiate overdose in Rhode Island, 1997-20021. eng, Subst Use Misuse 41, 1209–1226, (http://dx.doi.org/10.1080/10826080600751898) (2006).

3. Klimas, J., O’Reilly, M., Egan, M., Tobin, H., Bury, G., Urban overdose hotspots: a 12-month prospective study in Dublin ambulance services. eng, Am J Emerg Med 32, 1168–1173, (http://dx.doi.org/10.1016/j.ajem.2014.07.017) (2014).

4. Ryan, C. W., Spatial point pattern analysis of prehospital naloxone administrations (https://www.biorxiv.org/content/early/2017/01/16/100412).

5. Xu, J., Griffiths, E., Shooting on the street: measuring the spatial influence of physical features on gun violence in a bounded street network, Journal of quantitative criminology 33, 237–253 (2017).

6. Merikangas, K. R., McClair, V. L., Epidemiology of substance use disorders. Human genetics 131, 779–789, ISSN: 1432-1203 (6 2012).

7. Baddeley, A., Rubak, E., Turner, R., Spatial point patterns: methodology and applications with R (CRC Press, Taylor & Francis Group, Boca Raton London New York, 2016), ISBN: 9781482210200.

8. Hosmer, D., Lemeshow, S., Rodney, S., Applied logistic regression (Wiley, Hoboken, New Jersey, 2013), ISBN: 9780470582473.

9. Besag, J., Newell, J., The Detection of Clusters in Rare Diseases, Journal of the Royal Statistical Society. Series A (Statistics in Society) 154, 143–155, ISSN: 09641998, 1467985X, (http://www.jstor.org/stable/2982708) (1991).

10. Xu, G., Waagepetersen, R., Guan, Y., Stochastic Quasi-likelihood for Case-Control Point Pattern Data,. Journal of the American Statistical Association 0, 0–0, eprint: https://doi.org/10.1080/01621459.2017.1421543, (https://doi.org/10.1080/01621459.2017.1421543) (2018).

