## Supplementary material for "An inhomogeneous Poisson model of the spatial intensity of opioid overdoses, as represented by EMS naloxone use": R code used for anlaysis (with filename extension changed to .txt so as to work with BioRxiv upload)

### R code from vignette source 'e:/DATA/SCHOLAR/NaloxoneSpatialResearch/BioRxiv/BioRxiv2/NaloxoneSpatialInhomogeneousPoissonModel-BioRxiv-June2018.Rnw'

###################################################
### code chunk number 1: options
###################################################
setwd("E:/DATA/SCHOLAR/NaloxoneSpatialResearch/BioRxiv/BioRxiv2")
# setwd("I:/BCHD/Opioids") # change drive letter as needed
# setwd("F:/BCHD/Opioids") # change drive letter as needed
# setwd("/media/IronKey USB/BCHD/Opioids")
# setwd("/media/ryancwrf/Imation USB/BCHD/Opioids")
# setwd("G:/BCHD/Opioids")
# .libPaths( c( .libPaths(), "\\\\hdfp1/users/cwr080961/R/win-library/3.1") ) # work-around after I got new computer October 2014
options(SweaveSyntax="SweaveSyntaxNoweb")  # only needed at BCHD computer


###################################################
### code chunk number 2: spatialoptions
###################################################
## adjust the two lines below as needed by platform and operating system
# setwd("I:/BCHD/Opioids/NaloxoneSpatialAnalysis")
# setwd("F:/BCHD/Opioids/NaloxoneSpatialAnalysis")
# setwd("/media/ryancwrf/Imation USB/BCHD/Opioids/NaloxoneSpatialAnalysis")
# setwd("G:/BCHD/Opioids/NaloxoneSpatialAnalysis")
library(grDevices)
library(dplyr)
library(stringr)
library(Hmisc)
library(shapefiles)
library(spatstat)
library(maptools)
library(sp)
library(RColorBrewer)
library(rgdal)
library(tmap)
library(broom)
my.red.colors <- brewer.pal(n=7, "Reds")


###################################################
### code chunk number 3: somesetup
###################################################
## a function I will need
LRtests.f <- function(x, y){anova(x, y, test = "LR")[2,4]}  ## the [2,4] element is the p-value for the LR test
## create color ramps
my.uni.reds <- brewer.pal(7, "Reds")
my.div <- brewer.pal(11, "RdYlGn")
## number of replications/simulations
B <- 499
## a place to set the seed for the random number generator
set.seed(27)


###################################################
### code chunk number 4: wrangling
###################################################
nysArea <- readOGR("E:/DATA/GeographicData", "cty036")
nysAreaUTM <- spTransform(nysArea, CRS("+proj=utm +zone=18 +ellps=GRS80 +units=km +no_defs"))
sremsAreaUTM <- subset(nysAreaUTM, NAME=="Broome" | NAME=="Tioga" | NAME=="Chenango")
broomeAreaUTM <- subset(nysAreaUTM, NAME=="Broome")
tiogaAreaUTM <- subset(nysAreaUTM, NAME=="Tioga")
chenangoAreaUTM <- subset(nysAreaUTM, NAME=="Chenango")
temp.sremsWindow <- as(sremsAreaUTM, "owin")
temp.sremsWindow.2 <- closing(temp.sremsWindow, r=0.00001) # to collapse the 3-polygon window into a single window for the whole SREMS region
sremsWindow <- expand.owin(temp.sremsWindow.2, distance=0.001) # expand the window by 1 meter (0.001 km) all around, to avoid some road segments being considered outside the window
nysCensusTracts <- readOGR("E:/DATA/GeographicData/NYScensusTracts", "tl_2014_36_tract")
## keep the projection consistent for all objects
nysCensusTracts <- spTransform(nysCensusTracts, CRS("+proj=utm +zone=18 +ellps=GRS80 +units=km +no_defs"))
nysCensusTracts@data$geoid <- as.character(nysCensusTracts@data$GEOID)
## calculate surface area of each census tract in NYS, and add it as a variable
## to nysCensusTracts@data
p <- slot(nysCensusTracts, "polygons")
v <- lapply(p, function(z) { SpatialPolygons(list(z)) })
nysCensusTracts@data$surfaceArea <- sapply(v, area)

sremsCensusTracts <- subset(nysCensusTracts, COUNTYFP %in% c("007", "017", "107"))
## need the srems census tracts as a window, or pixel image, or tesselation for ppm() command
sremsCTwindow <- as(sremsCensusTracts, "owin")


## read in Household Poverty by census tract, which is simply a dataframe, nothing spatial
nysHouseholdPoverty <- read.csv("E:/DATA/GeographicData/NYSpovertyCensusTract/ACS_14_5YR_B17017.csv")
## select only the necessary variables
nysPoverty <- subset(nysHouseholdPoverty, select = c(GEO.id, HD01_VD01, HD01_VD02))
## create a character geoid variable for each census tract (CT)
nysPoverty$geoid <- str_sub(nysPoverty$GEO.id, start = 10)
## combine the poverty measurements with the spatial polygons dataframe of NYS CTs
temp.pov <-  merge(nysCensusTracts, nysPoverty, by = "geoid")
## calculate proportion of households in poverty, for each CT
temp.pov$propPoverty <- temp.pov$HD01_VD02/temp.pov$HD01_VD01
## discard all non-SREMS CTs
sremsPoverty <- subset(temp.pov, COUNTYFP %in% c("007", "017", "107"))

## now conduct similar operations on Housing Tenure
nysHousingTenure <- read.csv("E:/DATA/GeographicData/NYShousingTenureCensusTract/ACS_14_5YR_B25038.csv")
nysTenure <- subset(nysHousingTenure, select = c(GEO.id, HD01_VD01, HD01_VD02))
nysTenure$geoid <- str_sub(nysTenure$GEO.id, start = 10)
temp.ten <-  merge(nysCensusTracts, nysTenure, by = "geoid")
temp.ten$propOwnOcc <- temp.ten$HD01_VD02/temp.ten$HD01_VD01
sremsTenure <- subset(temp.ten, COUNTYFP %in% c("007", "017", "107"))

## now conduct similar operations on the
## total population and pop density, and
## PERCENT of people in age bracket 20 to 44
## PERCENT is provided DIRECTLY in Census Bureau files; I will change it to proportion
## for consistency with other variables
## HC01_EST_VC01 is total pop in the tract
nysAgeDistribution <- read.csv("E:/DATA/GeographicData/NYSAgeDistributionCensusTract/ACS_14_5YR_S0101.csv")
nysAgeDistribution <- subset(nysAgeDistribution, select = c(GEO.id,        ## geoid
                                                            HC01_EST_VC01, ## total population in tract
                                                            HC01_EST_VC07, ## percent age 20-24
                                                            HC01_EST_VC08, ## percent age 25-29
                                                            HC01_EST_VC09, ## percent age 30-34
                                                            HC01_EST_VC10, ## percent age 35-39
                                                            HC01_EST_VC11)) # percent age 40-44
nysAgeDistribution$totalPop <- nysAgeDistribution$HC01_EST_VC01
nysAgeDistribution$Age2044 <- rowSums(nysAgeDistribution[, 3:7])
nysAgeDistribution$geoid <- str_sub(nysAgeDistribution$GEO.id, start = 10)
temp.agedist <-  merge(nysCensusTracts, nysAgeDistribution, by  = "geoid")
temp.agedist$propAge2044 <- temp.agedist$Age2044/100  ## changes it to proportion
sremsAgeDistribution <- subset(temp.agedist, COUNTYFP %in% c("007", "017", "107"))
sremsAgeDistribution@data$popDensity <- sremsAgeDistribution@data$totalPop / sremsAgeDistribution@data$surfaceArea  ## people per sq km


## sremsCTdata <- merge(sremsAgeDistribution, sremsPoverty, by = "geoid")
## sremsCTdata2 <- merge(sremsCTdata, sremsTenure, by = "geoid", all = FALSE)


## read in point pattern data for cases and controls
cases <- read.table("E:/DATA/Scholar/NaloxoneSpatialResearch/NaloxoneSpatialCasesAnonymous-02-15-2016.txt")
controls <- read.table("E:/DATA/Scholar/NaloxoneSpatialResearch/NaloxoneSpatialControlsAnonymous-02-15-2016.txt")
cases.and.controls <- read.table("E:/DATA/Scholar/NaloxoneSpatialResearch/NaloxoneSpatialCasesAndControlsAnonymous-02-15-2016.txt")
## convert distances (coordinates) from meters to kilometers
## these lines will also delete the "marks" column, which will be added back later
cases <- cases[,c("x", "y")]/1000
controls <- controls[,c("x", "y")]/1000
cases.and.controls <- cases.and.controls[,c("x", "y")]/1000


## create point patterns inside the sremsWindow
cases.ppp <- ppp(cases$x, cases$y, window=sremsWindow)
unitname(cases.ppp) <- c("km")
marks(cases.ppp) <- "case"
controls.ppp <- ppp(controls$x, controls$y, window=sremsWindow)
unitname(controls.ppp) <- c("km")
marks(controls.ppp) <- "control"
cases.and.controls.ppp <- superimpose(controls.ppp, cases.ppp)
## the relrisk() function in spatstat (I might use it later, in some other analysis) will require factor type marks, with the first of the levels representing controls
cases.and.controls.ppp$marks <- factor(cases.and.controls.ppp$marks, levels=c("control", "case"))
# Regarding the issue of duplicate points:
# There are many. Multiplicities in among the controls rangs to 900+, which is absurd
# I think that is a consequence of some idiosyncrasies in the ImageTrend database.
# Surely there will be some locations that receive several EMS calls during the study
# period. But not 900.
# Upon reflection, I decided to eliminate duplicate points. The dataset then becomes:
# those locations to which EMS has resonded at least once during the study period, some with naloxone
# and some without. I think that is a reasonable and  defensible approach
unique.cases.ppp <- unique(cases.ppp)
unique.controls.ppp <- unique(controls.ppp)
unique.cases.and.controls.ppp <- unique(cases.and.controls.ppp)


srems.roads <- readOGR("E:/DATA/Scholar/NaloxoneSpatialResearch", "SREMSstreetlines") # it appears to be already in [+proj=utm +zone=18 +datum=NAD83 +units=km +no_defs +ellps=GRS80 +towgs84=0,0,0]
# but just to make sure
srems.roads.UTM <- spTransform(srems.roads, CRS("+proj=utm +zone=18 +ellps=GRS80 +units=km +no_defs"))
srems.roads.psp <- as.psp(srems.roads.UTM)
srems.roads.psp <- unmark(srems.roads.psp)

# the code below is needed because simply subsetting srems.roads.psp to sremsWindow via []
# is very memory intensive, and also seems to leave some stray segment endpoints outside the sremsWindow
marks(srems.roads.psp) <- inside.owin(srems.roads.psp$ends$x0, srems.roads.psp$ends$y0, sremsWindow) &  inside.owin(srems.roads.psp$ends$x1, srems.roads.psp$ends$y1, sremsWindow) # are both the starting point and ending point of every segment inside the window? Keep those that are.
srems.roads.psp.2 <- srems.roads.psp[marks(srems.roads.psp) == TRUE]
srems.roads.psp.2 <- unmark(srems.roads.psp.2)
srems.roads.psp <- srems.roads.psp.2
road.length.summary <- summary(srems.roads.psp)$len


## create density of all calls
## r now in km
radius <- seq(0,20, by=0.005)  # set r so that all of the G, F, and K functions from the simulations use the same values of r
dbw <- bw.diggle(unique.cases.and.controls.ppp)
pplbw <- bw.ppl(unique.cases.and.controls.ppp)
all.calls.density <- density(unique.cases.and.controls.ppp, bw=pplbw, adjust=0.2, positive=TRUE, dimyx=c(512,512))
sampling.dens <- all.calls.density

########################################################################
#### Add old code that I used for initial descriptive, non-model-based clustering analysis at CSTE 2017 Boise. This is for Upstat 2018 in Rochester.

## SREMS roads map
pdf(file="sremsRoads.pdf")
par(mar=rep(1, 4), oma = rep(1,4))
plot(srems.roads.psp, main="")
dev.off()

## SREMS orientation map
sremsCountyNames <- c("Broome", "Tioga", "Chenango")
pdf(file = "SREMSorientationMap.pdf", width = 7)
plot(nysAreaUTM[nysAreaUTM$NAME %in% sremsCountyNames], col = nysAreaUTM$NAME %in% sremsCountyNames)
dev.off()

## Overview plot
## remember things are in km now, not meters
ys <- yardstick(x0=377.999, y0=467.4000, x1=387.999, y1=4674.000, txt="10 km", win=sremsWindow)
pdf(file="NaloxoneOverviewPlot.pdf")
par(mar=rep(1, 4), oma = rep(1,4))
plot(rjitter(unique.cases.and.controls.ppp, radius=0.5),  pch=c(1,16), cex=c(0.6,0.6), cols=c("blue", "red"), main="")
plot(ys, angle=90, cex=0.8, lwd=2)
dev.off()


###################################################
### code chunk number 5: preparetocheckclustering
###################################################
# will need these for all G, F, and K, to create and plot simulation envelopes
# this will find 95% point-wise acceptance bands (with a grain of salt)
LB.function <- function(x){quantile(x, probs=0.025, na.rm=TRUE)}
UB.function <- function(x){quantile(x, probs=0.975, na.rm=TRUE)}

ratio <- unique.cases.ppp$n/unique.cases.and.controls.ppp$n
N <- unique.cases.ppp$n
BB <- 250 ## number of simulations for my original, non-model-based analysis


###################################################
### code chunk number 6: summariesfromobservednaloxoneincidents
###################################################
# get the summary functions for the observed cases
G.observed.cases <- Gest(unique.cases.ppp, r=radius)
G.observed.cases.values <- G.observed.cases$rs
F.observed.cases <- Fest(unique.cases.ppp, r=radius)
F.observed.cases.values <- F.observed.cases$rs
K.observed.cases <- Kinhom(unique.cases.ppp, r=radius, correction="translate")
K.observed.cases.values <- K.observed.cases$trans
L.observed.cases.values <- sqrt(K.observed.cases.values/pi)
ipcf.observed.cases <- pcfinhom(unique.cases.ppp,  r = radius, normpower = 2, renormalize = TRUE, leaveoneout = TRUE)
ipcf.observed.cases.values <- ipcf.observed.cases$trans


###################################################
### code chunk number 7: simulations1
###################################################
# Monte Carlo simulations from the observed locations of all actual incidents
B.1 <- BB  # number of simulations. Change as needed
G.values.1 <- matrix(NA, nrow=length(radius), ncol=B.1)
F.values.1 <- matrix(NA, nrow=length(radius), ncol=B.1)
K.values.1 <- matrix(NA, nrow=length(radius), ncol=B.1)
ipcf.values.1 <- matrix(NA, nrow=length(radius), ncol=B.1)
example.samples.1 <- list(rep(NA, 4))
for (i in 1:B.1){
    ## this will extract a random sample of all incident locations,
    ## equal in size to the number of naloxone cases
    keep <- sample(1:npoints(unique.cases.and.controls.ppp), size = N, replace = FALSE)
    temp.ppp.2  <- unmark(unique.cases.and.controls.ppp[keep])
    temp.ppp <- temp.ppp.2  ## for convenience with legacy code
    if (i <= 4) {example.samples.1[[i]] <- temp.ppp}
    # d <- 6/sqrt(2) # to move each point 6 meters, perpendicularly, from its road segment
    # temp.ppp <- shift(temp.ppp.1, vec=c(d,d))
    # temp.ppp <- superimpose(rjitter(subset(temp.ppp.1, marks=="B"), radius=6), subset(temp.ppp.1, marks=="A"))
    # save one unadjusted 183-point sample, for illustrative purposes outside the loop
    # save one corrected illustrative 183-point sample
    G.values.1[,i] <- Gest(temp.ppp, r=radius)$rs  # uses the reduced sample or "border" method.  There are other options
    F.values.1[,i] <- Fest(temp.ppp, r=radius)$rs  # uses the reduced sample or "border" method.  There are other options
    K.values.1[,i] <- Kinhom(temp.ppp, r=radius, correction="translate")$trans
    ipcf.values.1[,i] <- pcfinhom(temp.ppp, r = radius, normpower = 2, renormalize = TRUE, leaveoneout = TRUE)$trans
 }
# and get L simply from K
L.values.1 <- sqrt(K.values.1/pi)
# collect the examples of simulations into a solist
example.samples.1 <- as.solist(example.samples.1)


###################################################
### code chunk number 8: examplesamples1
###################################################
pdf(file="ExampleSamples1.pdf",width=8)
par(mar=rep(1, 4), oma = rep(1,4))
plot(example.samples.1, nrows=2, ncols=2, main="", main.panel="", arrange=TRUE, equal.scales=TRUE, mar.panel=1, par(mar=rep(0.5, 4)))
dev.off()


###################################################
### code chunk number 9: keyplots
###################################################
# get minimums, maximums, and 95% upper and lower bounds of
# the sampling distributions
sim.G.min.1 <- apply(G.values.1, 1, FUN=min)
sim.G.max.1 <- apply(G.values.1, 1, FUN=max)
sim.G.LB.1 <- apply(G.values.1, 1, FUN=LB.function)
sim.G.UB.1 <- apply(G.values.1, 1, FUN=UB.function)
sim.F.min.1 <- apply(F.values.1, 1, FUN=min)
sim.F.max.1 <- apply(F.values.1, 1, FUN=max)
sim.F.LB.1 <- apply(F.values.1, 1, FUN=LB.function)
sim.F.UB.1 <- apply(F.values.1, 1, FUN=UB.function)
sim.K.min.1 <- apply(K.values.1, 1, FUN=min)
sim.K.max.1 <- apply(K.values.1, 1, FUN=max)
sim.K.LB.1 <- apply(K.values.1, 1, FUN=LB.function)
sim.K.UB.1 <- apply(K.values.1, 1, FUN=UB.function)
sim.L.LB.1 <- apply(L.values.1, 1, FUN=LB.function)
sim.L.UB.1 <- apply(L.values.1, 1, FUN=UB.function)
sim.L.min.1 <- apply(L.values.1, 1, FUN=min)
sim.L.max.1 <- apply(L.values.1, 1, FUN=max)
sim.ipcf.LB.1 <- apply(ipcf.values.1, 1, FUN=LB.function)
sim.ipcf.UB.1 <- apply(ipcf.values.1, 1, FUN=UB.function)
sim.ipcf.min.1 <- apply(ipcf.values.1, 1, FUN=min)
sim.ipcf.max.1 <- apply(ipcf.values.1, 1, FUN=max)
## make plots

## first graph: G
pdf(file = "G1.pdf", width = 7)
par(mgp = c(2, 1, 0))
plot(sim.G.UB.1 ~ radius, type = "n", ylab = "", xlab = "distance in kilometers", xlim = c(0,10))
lines(sim.G.LB.1 ~ radius, type = "n")
polygon(c(radius, rev(radius)), c(sim.G.UB.1, rev(sim.G.LB.1)), col = "grey")
lines(G.observed.cases.values ~ radius, type = "l", lwd = 2, col = "red")
lines(sim.G.max.1 ~ radius, type = "l", lty = 2, col = "blue")
lines(sim.G.min.1 ~ radius, type = "l", lty = 2, col = "blue")
legend(legend=c("G function", "observed cases", "95% pointwise simulation envelope", "minimum and maximum simulation"), lty = c(NA, 1,1,2), lwd = c(NA,2,8,1), col = c("white", "red", "grey", "blue"), "bottomright")
dev.off()
## second graph: F
pdf(file = "F1.pdf", width = 7)
par(mgp = c(2, 1, 0))
plot(sim.F.UB.1 ~ radius, type = "n", ylab = "", xlab = "distance in kilometers")
lines(sim.F.LB.1 ~ radius, type = "n")
polygon(c(radius, rev(radius)), c(sim.F.UB.1, rev(sim.F.LB.1)), col = "grey")
lines(F.observed.cases.values ~ radius, type = "l", lwd = 2, col = "red")
lines(sim.F.max.1 ~ radius, type = "l", lty = 2, col = "blue")
lines(sim.F.min.1 ~ radius, type = "l", lty = 2, col = "blue")
legend(legend=c("F function", "observed cases", "95% pointwise simulation envelope", "minimum and maximum simulation"), lty = c(NA, 1,1,2), lwd = c(NA,2,8,1), col = c("white", "red", "grey", "blue"), "bottomright")
dev.off()
##third graph: K
pdf(file = "K1.pdf", width = 7)
par(mgp = c(2, 1, 0))
plot(sim.K.UB.1 ~ radius, type = "n", ylab = "", xlab = "distance in kilometers")
lines(sim.K.LB.1 ~ radius, type = "n")
polygon(c(radius, rev(radius)), c(sim.K.UB.1, rev(sim.K.LB.1)), col = "grey")
lines(K.observed.cases.values ~ radius, type = "l", lwd = 2, col = "red")
lines(sim.K.max.1 ~ radius, type = "l", lty = 2, col = "blue")
lines(sim.K.min.1 ~ radius, type = "l", lty = 2, col = "blue")
legend(legend=c("inhomogeneous K function", "observed cases", "95% pointwise simulation envelope", "minimum and maximum simulation"), lty = c(NA, 1,1,2), lwd = c(NA,4,20,2), col = c("white", "red", "grey", "blue"), "topleft", cex = 1.3)
arrows(1,350,3,220, col = "purple", lwd = 6)
dev.off()
## fourth graph: L
pdf(file = "L1.pdf", width = 7)
par(mgp = c(2, 1, 0))
plot(sim.L.UB.1 ~ radius, type = "n", ylab = "", xlab = "distance in kilometers")
lines(sim.L.LB.1 ~ radius, type = "n")
polygon(c(radius, rev(radius)), c(sim.L.UB.1, rev(sim.L.LB.1)), col = "grey")
lines(L.observed.cases.values ~ radius, type = "l", lwd = 2, col = "red")
lines(sim.L.max.1 ~ radius, type = "l", lty = 2, col = "blue")
lines(sim.L.min.1 ~ radius, type = "l", lty = 2, col = "blue")
legend(legend=c("L function", "observed cases", "95% pointwise simulation envelope", "minimum and maximum simulation"), lty = c(NA, 1,1,2), lwd = c(NA,2,8,1), col = c("white", "red", "grey", "blue"), "bottomright")
dev.off()
## fifth graph: ipcf
pdf(file = "ipcf1.pdf", width = 7)
par(mgp = c(2, 1, 0))
## I'm making artistic choices of xlim and ylim here, to maximize info:ink ratio
## draw a gray simulation envelope for the inhompcf ratios under null hypothesis
## draw the line representing the observed cases ipcf last, so it will not be "under"
## the gray simulation envelope
plot(sim.ipcf.UB.1 ~ radius, type = "n", ylim = c(0,100), xlim = c(0,3), ylab = "", xlab = "distance in kilometers")
lines(sim.ipcf.LB.1 ~ radius, type = "n")
polygon(c(radius, rev(radius)), c(sim.ipcf.UB.1, rev(sim.ipcf.LB.1)), col = "grey")
lines(ipcf.observed.cases.values ~ radius, type = "l", lwd = 2, col = "red")
lines(sim.ipcf.max.1 ~ radius, type = "l", lty = 2, col = "blue")
lines(sim.ipcf.min.1 ~ radius, type = "l", lty = 2, col = "blue")
legend(legend=c("inhomogeneous pair correlation function", "observed cases", "95% pointwise simulation envelope", "minimum and maximum simulation"), lty = c(NA, 1,1,2), lwd = c(NA,2,8,1), col = c("white", "red", "grey", "blue"), "topright")
dev.off()


###################################################
### code chunk number 10: simulations.2
###################################################
# Monte Carlo simulation from 2-D density derived from locations of all incidents
B.2 <- BB  # number of simulations. Change as needed
G.values.2 <- matrix(NA, nrow=length(radius), ncol=B.2)
F.values.2 <- matrix(NA, nrow=length(radius), ncol=B.2)
K.values.2 <- matrix(NA, nrow=length(radius), ncol=B.2)
ipcf.values.2 <- matrix(NA, nrow=length(radius), ncol=B.2)
proj.distances <- matrix(NA, nrow=N, ncol=B.2)
example.samples.2 <- list(rep(NA, 4))
for (i in 1:B.2){
    temp.ppp.2  <- rpoint(n=N, f=sampling.dens)
    # adjust the resulting simulated point pattern
    temp.ppp.2b <- project2segment(temp.ppp.2, srems.roads.psp)
    proj.distances[,i] <- temp.ppp.2b$d
    temp.ppp <- temp.ppp.2b$Xproj
    if (i <= 4) {example.samples.2[[i]] <- temp.ppp}
    # d <- 6/sqrt(2) # to move each point 6 meters, perpendicularly, from its road segment
    # temp.ppp <- shift(temp.ppp.1, vec=c(d,d))
    # temp.ppp <- superimpose(rjitter(subset(temp.ppp.1, marks=="B"), radius=6), subset(temp.ppp.1, marks=="A"))
    # save one unadjusted 183-point sample, for illustrative purposes outside the loop
    # save one corrected illustrative 183-point sample
    G.values.2[,i] <- Gest(temp.ppp, r=radius)$rs  # uses the reduced sample or "border" method.  There are other options
    F.values.2[,i] <- Fest(temp.ppp, r=radius)$rs  # uses the reduced sample or "border" method.  There are other options
    K.values.2[,i] <- Kinhom(temp.ppp, r=radius, correction="translate")$trans
     ipcf.values.2[,i] <- pcfinhom(temp.ppp, r = radius, normpower = 2, renormalize = TRUE, leaveoneout = TRUE)$trans
 }
# and get L simply from K
L.values.2 <- sqrt(K.values.2/pi)
# collect the examples of simulations into a solist
example.samples.2 <- as.solist(example.samples.2)


###################################################
### code chunk number 11: howdidadjustmentwork
###################################################
proj.distance.summary <- fivenum(proj.distances)


###################################################
### code chunk number 12: examplesamples
###################################################
pdf(file="ExampleSamples2.pdf",width=8)
plot(example.samples.2, fg = "white", nrows=2, ncols=2, main="", main.panel="", arrange=TRUE, equal.scales=TRUE, mar.panel=1, par(mar=rep(0.5, 4)))
dev.off()


###################################################
### code chunk number 13: keyplots
###################################################
# get minimums, maximums, and 95% upper and lower bounds of
# the sampling distributions
sim.G.min.2 <- apply(G.values.2, 1, FUN=min)
sim.G.max.2 <- apply(G.values.2, 1, FUN=max)
sim.G.LB.2 <- apply(G.values.2, 1, FUN=LB.function)
sim.G.UB.2 <- apply(G.values.2, 1, FUN=UB.function)
sim.F.min.2 <- apply(F.values.2, 1, FUN=min)
sim.F.max.2 <- apply(F.values.2, 1, FUN=max)
sim.F.LB.2 <- apply(F.values.2, 1, FUN=LB.function)
sim.F.UB.2 <- apply(F.values.2, 1, FUN=UB.function)
sim.K.min.2 <- apply(K.values.2, 1, FUN=min)
sim.K.max.2 <- apply(K.values.2, 1, FUN=max)
sim.K.LB.2 <- apply(K.values.2, 1, FUN=LB.function)
sim.K.UB.2 <- apply(K.values.2, 1, FUN=UB.function)
sim.L.LB.2 <- apply(L.values.2, 1, FUN=LB.function)
sim.L.UB.2 <- apply(L.values.2, 1, FUN=UB.function)
sim.L.min.2 <- apply(L.values.2, 1, FUN=min)
sim.L.max.2 <- apply(L.values.2, 1, FUN=max)
sim.ipcf.LB.2 <- apply(ipcf.values.2, 1, FUN=LB.function)
sim.ipcf.UB.2 <- apply(ipcf.values.2, 1, FUN=UB.function)
sim.ipcf.min.2 <- apply(ipcf.values.2, 1, FUN=min)
sim.ipcf.max.2 <- apply(ipcf.values.2, 1, FUN=max)
## make plots


## first graph: G2
pdf(file = "G2.pdf", width = 7)
par(mgp = c(2, 1, 0))
plot(sim.G.UB.2 ~ radius, type = "n", ylab = "", xlab = "distance in kilometers", xlim = c(0,10))
lines(sim.G.LB.2 ~ radius, type = "n")
polygon(c(radius, rev(radius)), c(sim.G.UB.2, rev(sim.G.LB.2)), col = "grey")
lines(G.observed.cases.values ~ radius, type = "l", lwd = 2, col = "red")
lines(sim.G.max.2 ~ radius, type = "l", lty = 2, col = "blue")
lines(sim.G.min.2 ~ radius, type = "l", lty = 2, col = "blue")
legend(legend=c("G function", "observed cases", "95% pointwise simulation envelope", "minimum and maximum simulation"), lty = c(NA, 1,1,2), lwd = c(NA,2,8,1), col = c("white", "red", "grey", "blue"), "bottomright")
dev.off()
## second graph: F2
pdf(file = "F2.pdf", width = 7)
par(mgp = c(2, 1, 0))
plot(sim.F.UB.2 ~ radius, type = "n", ylab = "", xlab = "distance in kilometers")
lines(sim.F.LB.2 ~ radius, type = "n")
polygon(c(radius, rev(radius)), c(sim.F.UB.2, rev(sim.F.LB.2)), col = "grey")
lines(F.observed.cases.values ~ radius, type = "l", lwd = 2, col = "red")
lines(sim.F.max.2 ~ radius, type = "l", lty = 2, col = "blue")
lines(sim.F.min.2 ~ radius, type = "l", lty = 2, col = "blue")
legend(legend=c("F function", "observed cases", "95% pointwise simulation envelope", "minimum and maximum simulation"), lty = c(NA, 1,1,2), lwd = c(NA,2,8,1), col = c("white", "red", "grey", "blue"), "bottomright")
dev.off()
##third graph: K2
pdf(file = "K2.pdf", width = 7)
par(mgp = c(2, 1, 0))
plot(sim.K.UB.2 ~ radius, type = "n", ylab = "", xlab = "distance in kilometers")
lines(sim.K.LB.2 ~ radius, type = "n")
polygon(c(radius, rev(radius)), c(sim.K.UB.2, rev(sim.K.LB.2)), col = "grey")
lines(K.observed.cases.values ~ radius, type = "l", lwd = 2, col = "red")
lines(sim.K.max.2 ~ radius, type = "l", lty = 2, col = "blue")
lines(sim.K.min.2 ~ radius, type = "l", lty = 2, col = "blue")
legend(legend=c("inhomogeneous K function", "observed cases", "95% pointwise simulation envelope", "minimum and maximum simulation"), lty = c(NA, 1,1,2), lwd = c(NA,2,8,1), col = c("white", "red", "grey", "blue"), "topleft")
dev.off()
## fourth graph: L2
pdf(file = "L2.pdf", width = 7)
par(mgp = c(2, 1, 0))
plot(sim.L.UB.2 ~ radius, type = "n", ylab = "", xlab = "distance in kilometers")
lines(sim.L.LB.2 ~ radius, type = "n")
polygon(c(radius, rev(radius)), c(sim.L.UB.2, rev(sim.L.LB.2)), col = "grey")
lines(L.observed.cases.values ~ radius, type = "l", lwd = 2, col = "red")
lines(sim.L.max.2 ~ radius, type = "l", lty = 2, col = "blue")
lines(sim.L.min.2 ~ radius, type = "l", lty = 2, col = "blue")
legend(legend=c("L function", "observed cases", "95% pointwise simulation envelope", "minimum and maximum simulation"), lty = c(NA, 1,1,2), lwd = c(NA,2,8,1), col = c("white", "red", "grey", "blue"), "bottomright")
dev.off()


## fifth graph: ipcf2
pdf(file = "ipcf2.pdf", width = 7)
par(mgp = c(2, 1, 0))
## I'm making artistic choices of xlim and ylim here, to maximize info:ink ratio
## draw a gray simulation envelope for the inhompcf ratios under null hypothesis
## draw the line representing the observed cases ipcf last, so it will not be "under"
## the gray simulation envelope
plot(sim.ipcf.UB.2 ~ radius, type = "n", ylim = c(0,100), xlim = c(0,3), ylab = "", xlab = "distance in kilometers")
lines(sim.ipcf.LB.2 ~ radius, type = "n")
polygon(c(radius, rev(radius)), c(sim.ipcf.UB.2, rev(sim.ipcf.LB.2)), col = "grey")
lines(ipcf.observed.cases.values ~ radius, type = "l", lwd = 2, col = "red")
lines(sim.ipcf.max.2 ~ radius, type = "l", lty = 2, col = "blue")
lines(sim.ipcf.min.2 ~ radius, type = "l", lty = 2, col = "blue")
legend(legend=c("inhomogeneous pair correlation function", "observed cases", "95% pointwise simulation envelope", "minimum and maximum simulation"), lty = c(NA, 1,1,2), lwd = c(NA,2,8,1), col = c("white", "red", "grey", "blue"), "topright")
dev.off()


###################################################
### code chunk number 14: preparepredictors
###################################################
## for modeling with ppm(), I have to get the spatialpolygonsdataframes for poverty, housing tenure,
## and age distribution into forms that are acceptable inputs for predictors in ppm(). SPDFs are not.
## I will turn them into functions as in f(x,y), functions of the x and y coordinates, and thence into images.
## The problem with funxy themselves is that they seem to pertain to the entire enclosing rectangle, with NA
## values in all pixels outside the sremsWindow. These NA values cause downstream problems with calculating
## residuals for model diagnosits. I will remedy this with the intersect.owin command, in the chunk labeled fixwindowproblem, further below.
## First make a tessalation, with each census tract a tile
## then turn the tesselation into a funxy
## and then into a pixel image, restricted to the sremsWindow

### Poverty
p <- slot(sremsPoverty, "polygons")
v <- lapply(p, function(z) { SpatialPolygons(list(z)) })
sat <- tess(tiles = lapply(v, as.owin) )
pov.f <- as.function.tess(sat, values = sremsPoverty@data$propPoverty)

### Housing tenure
p <- slot(sremsTenure, "polygons")
v <- lapply(p, function(z) { SpatialPolygons(list(z)) })
sat <- tess(tiles = lapply(v, as.owin))
ten.f <- as.function.tess(sat, values = sremsTenure@data$propOwnOcc)

### proportion of residents between 20 and 44
p <- slot(sremsAgeDistribution, "polygons")
v <- lapply(p, function(z) { SpatialPolygons(list(z)) })
sat <- tess(tiles = lapply(v, as.owin))
age2044.f <- as.function.tess(sat, values = sremsAgeDistribution@data$propAge2044)

### population density
p <- slot(sremsAgeDistribution, "polygons")
v <- lapply(p, function(z) { SpatialPolygons(list(z)) })
sat <- tess(tiles = lapply(v, as.owin))
popDens.f <- as.function.tess(sat, values = sremsAgeDistribution@data$popDensity)


## After creating some ppm models, I can simulate from them and calculate confidence bands for
## summary functions of clustering, such as F, G, Kinhom, L, pcfinhom, and such.
## I could use the envelope() function with Kest, Lest, etc, but that leaves the problem of the actual
## incident locations all being recorded as street addresses, so that the simulated locations would by
## definition be more diffuse (less clustered)
## So better to use the algorithm I used for my earlier work, such as in CSTE-Boise, in which I simulate
## just one ppp at a time, using simulate(m5, nsim = 1) and then displace the resulting points to the
## nearest road segment in srems.roads.psp, with project2segment(). The latter command will put the displaced
## point pattern into objectname$Xproj


## wrangle the convenience store locations

BroomeMiniMarts <- readOGR("E:/DATA/Scholar/NaloxoneSpatialResearch/ChengbinMiniMarts/Gas_Stations", "GasStation")
BroomeMiniMartsUTM <- spTransform(BroomeMiniMarts, CRS("+proj=utm +zone=18 +ellps=GRS80 +units=km +no_defs"))
BroomeMiniMarts.ppp <- as(BroomeMiniMartsUTM, "ppp")
BroomeMiniMarts.ppp <- unmark(BroomeMiniMarts.ppp)
Window(BroomeMiniMarts.ppp) <- Window(unique.cases.ppp)
## Tioga County
TiogaMiniMarts <- readOGR("E:/DATA/Scholar/NaloxoneSpatialResearch/ChengbinMiniMarts/Gas_Stations", "GasStation_Tioga")
TiogaMiniMartsUTM <- spTransform(TiogaMiniMarts, CRS("+proj=utm +zone=18 +ellps=GRS80 +units=km +no_defs"))
TiogaMiniMarts.ppp <- as(TiogaMiniMartsUTM, "ppp")
TiogaMiniMarts.ppp <- unmark(TiogaMiniMarts.ppp)
Window(TiogaMiniMarts.ppp) <- Window(unique.cases.ppp)
## ChenangoCounty
ChenangoMiniMarts <- readOGR("E:/DATA/Scholar/NaloxoneSpatialResearch/ChengbinMiniMarts/Gas_Stations", "GasStation_Chenango")
ChenangoMiniMartsUTM <- spTransform(ChenangoMiniMarts, CRS("+proj=utm +zone=18 +ellps=GRS80 +units=km +no_defs"))
ChenangoMiniMarts.ppp <- as(ChenangoMiniMartsUTM, "ppp")
ChenangoMiniMarts.ppp <- unmark(ChenangoMiniMarts.ppp)
Window(ChenangoMiniMarts.ppp) <- Window(unique.cases.ppp)
## Combine and isolate unique locations
MiniMarts <- unique(superimpose(BroomeMiniMarts.ppp, TiogaMiniMarts.ppp, ChenangoMiniMarts.ppp))

## a ppp object cannot be used as a predictor in ppm().
## I think the best approach would be to create a function representing the distance
## to the nearest MiniMart
MMdist <- distfun(MiniMarts)


###################################################
### code chunk number 15: descriptives
###################################################
ys <- yardstick(x0=377.999, y0=4674.000, x1=387.999, y1=4674.000, txt="10 km", win=sremsWindow)
pdf(file="NaloxoneOverviewPlot.pdf")
par(mar=rep(1, 4), oma = rep(1,4))
plot(rjitter(unique.cases.and.controls.ppp, radius=0.5),  pch=c(1,16), cex=c(0.6,0.6), cols=c("blue", "red"), main="")
plot(ys, angle=90, cex=0.8, lwd=2)
dev.off()
pdf(file = "popDensmap.pdf")
plot(popDens.f, col = my.uni.reds, main = "Population density")
dev.off()
pdf(file = "popDensdes.pdf")
plot(hist(sremsAgeDistribution@data$popDens), col = "grey", main = "Population density for 77 census tracts", xlab = "population density")
dev.off()
pdf(file = "povmap.pdf")
plot(pov.f, col = my.uni.reds, main = "Proportion of households in poverty")
dev.off()
pdf(file = "povdes.pdf")
plot(hist(sremsPoverty@data$propPov), col = "grey", main = "Proportion of households in poverty for 77 census tracts", xlab = "proportion below federal poverty level")
dev.off()
pdf(file = "tenmap.pdf")
plot(ten.f, col = my.uni.reds, main = "Proportion of owner-occupied residences")
dev.off()
pdf(file = "tendes.pdf")
plot(hist(sremsTenure@data$propOwnOcc), col = "grey", main = "Proportion of residences that are owner-occupied", xlab = "proportion owner-occupied")
dev.off()
pdf(file = "age2044map.pdf")
plot(age2044.f, col = my.uni.reds, main = "Proportion of people between ages 20 and 44")
dev.off()
pdf(file = "age2044des.pdf")
plot(hist(sremsAgeDistribution@data$propAge2044), col = "grey", main = "Proportion of people between ages 20 and 44, for 77 census tracts", xlab = "proportion between 20 and 44 years of age")
dev.off()
pdf(file = "MMdistmap.pdf")
plot(MMdist, col = my.uni.reds, main = "distance in km to nearest minimart")
plot(MiniMarts, col = "blue", pch = 16, add = TRUE)
dev.off()
## a histogram or density plot of MMdist itself doesn't make much sense
## but this may be insightful: distance from each case to nearest MM,
## versus distance from each control to nearest MM
pdf(file = "nncross.pdf")
plot(density(nncross(unique.cases.ppp, MiniMarts)$dist), col = "red", xlim = c(0,5), lwd = 2, main = "distribution of distances to nearest minimart", xlab = "distance (km) to nearest minimart")
lines(density(nncross(unique.controls.ppp, MiniMarts)$dist), lwd = 2)
legend("topright", legend = c("cases", "controls"), lwd = 2, col = c("red", "black"), cex = 1.5)
dev.off()


###################################################
### code chunk number 16: fixwindowproblem
###################################################
##  I'll use ten.f as the "archetypal" window
WW <- intersect.owin(Window(unique.cases.ppp), Window(ten.f))
unique.cases.ppp.2 <- unique.cases.ppp[WW]


###################################################
### code chunk number 17: univariatemodeling
###################################################
## univariate models
## consider m0 to be the ``null model''
m0 <-  ppm(unmark(unique.cases.ppp.2) ~ offset(log(all.calls.density)))
## all update() commands below are using fixdummy = TRUE by default ("highly recommended" per the help pages)
ma <- update(m0, ~ . + popDens.f)
mb <- update(m0, ~ . + pov.f)
mc <- update(m0, ~ . + ten.f)
md <- update(m0, ~ . + age2044.f)
me <- update(m0, ~ . + MMdist)

## univariate LR tests (better than the univariate Wald tests in summary.ppm()

LRtest.m0.f <- function(x){anova(m0, x, test = "LR")[2,4]}  ## the [2,4] element is the p-value for the LR test
univariateLRpvalues <- sapply(list(ma, mb, mc, md, me), FUN = LRtest.m0.f)  ## this works quite well, and no loop!

## all predictors are very signficantly different from 0 in univariate LR testing


###################################################
### code chunk number 18: multivariatemodeling
###################################################
## Multivariate model
## all update() commands below are using fixdummy = TRUE by default ("highly recommended" per the help pages)
## I chose to use unnecessarily verbose syntax to make it clearer what I am doing, and to support the model names
m1 <- update(m0, ~ . + popDens.f)
m2 <- update(m0, ~ . + popDens.f + pov.f)
m3 <- update(m0, ~ . + popDens.f + pov.f + ten.f)
m4 <- update(m0, ~ . + popDens.f + pov.f + ten.f + age2044.f)
m5 <- update(m0, ~ . + popDens.f + pov.f + ten.f + age2044.f + MMdist)


## Try dropping popDens.f,  pov.f, MMdist, and age2044.f one at a time
## all update() commands below are using fixdummy = TRUE by default ("highly recommended" per the help pages)
m6 <- update(m5, ~ . - popDens.f)
m7 <- update(m5, ~ . - pov.f)
m8 <- update(m5, ~ . - ten.f)
m9 <- update(m5, ~ . - age2044.f)
m10 <- update(m5, ~ . - MMdist)
##  LR testing of each predictor in m5 is a bit different from the Wald testing--and preferred
LRtests.m5 <- sapply(list(m6, m7, m8, m9, m10), m5, FUN = LRtests.f)  ## the output here is the LR test p-value for each term; NOT the coefficient on that term
extract.term.names.m5.f <- function(x){names(coef(m5))[which(!(names(coef(m5)) %in% names(coef(x))))]}
names(LRtests.m5) <- sapply(list(m6, m7, m8, m9, m10), FUN = extract.term.names.m5.f)

## get a nice table of LR signficance of each predictor in m5,
## along with the AICs if each is eliminated
## the drop1 command with the "Chisq" test will produce the LR p-values easily
singleTermDrop1table.m5 <- drop1(m5, test = "Chisq")

## including only the LR-test "significant" predictors (at alpha = 0.05) yields, essentially, mc: the model with ten.f as the only predictor:
m11 <- update(m5, ~ . - popDens.f - pov.f - age2044.f - MMdist)


## assess change in coefficient on ten.f (delta Beta hat percent in Hosmer and Lemeshow Applied Logistic Regression)
## use the broom package and dplyr to make human-readable dataframes

m5m6 <- left_join(tidy(coef(m5)), tidy(coef(m6)), by = c("names"))
m5m6m7 <- left_join(m5m6, tidy(coef(m7)), by = c("names"))
m5m6m7m8 <- left_join(m5m6m7, tidy(coef(m8)), by = "names")
m5m6m7m8m9 <- left_join(m5m6m7m8, tidy(coef(m9)), by = "names")
m5m6m7m8m9m10 <- left_join(m5m6m7m8m9, tidy(coef(m10)), by = "names")
m5m6m7m8m9m10m11 <- left_join(m5m6m7m8m9m10, tidy(coef(m11)), by = "names")
m.all <- m5m6m7m8m9m10m11
names(m.all) <- c("term", "m5", "m6", "m7", "m8", "m9", "m10", "m11")

## delta beta hat
## the definition of delta beta hat I use in the algorithm below is
## "reversed" from that used by Hosmer and Lemeshow in their classic text
## on applied logistic regression. Reversed in sign. I think it will
## make more sense to my audience this way than H&L's way.
delta.betahat.m5.f <- function(x){( (x - coef(m5)) / coef(m5) ) * 100}
percent.delta.betahat <- round(apply(m.all[ , -1], 2, FUN = delta.betahat.m5.f), 1) ## first column of m.all is rownames

## it looks like pov.f is the only predictor that is "expendable," in terms of both non-significant P-value AND changes in the other coefficients.
## So I will reinsert all but pov.f---popDens.f, age2044.f, and MMdist---even though they are not conventionally "significant", to make model 12
m12 <- m7  ## m7 is m5 with pov.f removed

## and check LR signficance of each predictor in m12
## the drop1 command with the "Chisq" test will produce the LR p-values easily
singleTermDrop1table.m12 <- drop1(m12, test = "Chisq")


###################################################
### code chunk number 19: m12diagnostics
###################################################
#### diagnostics for m12 Poisson model ####

## residuals
pres12 <- residuals(m12, "pearson")  ## pearson
res12 <- residuals(m12) ## raw
## notional cutoff for "significant" pearson residuals at 0.05 level
pearsonCutoff12 <- 2 * diagnose.ppm(m12, type = "Pearson", plot.it = FALSE)$smooth$sdp
pdf(file = "SmoothedPearsonResidualPlot-m12.pdf", width = 12)
plot(Smooth(pres12), col = my.div, main = "Smoothed Pearson residual plot")
plot(sat, add = TRUE)  ## sat is the latest value of the tesselation of sremsWindow, back from the wrangling stage
dev.off()


pres0 <- residuals(m0, "pearson")  ## pearson
res0 <- residuals(m0) ## raw
## notional cutoff for "significant" pearson residuals at 0.05 level
pearsonCutoff0 <- 2 * diagnose.ppm(m0, type = "Pearson", plot.it = FALSE)$smooth$sdp
pdf(file = "SmoothedPearsonResidualPlot-m0.pdf")
plot(Smooth(pres0), col = my.div, main = "Smoothed Pearson residual field for null model \n  (offset of all-calls density as the only predictor)")
plot(sat, add = TRUE)  ## sat is the latest value of the tesselation of sremsWindow, back from the wrangling stage
dev.off()


###################################################
### code chunk number 20: resulttables
###################################################
m12OR <- exp(coef(m12))
m12ORCI <- exp(confint(m12))
## recall and use percent.delta.betahat here
## recall and use singleTermDrop1table.m12 here too
model.table <- round(cbind(m12OR, m12ORCI), 3)


###################################################
### code chunk number 21: assessclusteringsimulations
###################################################
## assess how well the Poisson m12 accounts for the previously-observed clustering
## use a sample-generating expression that samples from the entire density of all calls, and displaces each
## sampled point to the nearest road
## based on Kinhom
## Note this explanation of global confidence/simulation/critical bands from R help for spatstat:
## "if global=TRUE, then the envelopes are determined as follows. First we calculate the theoretical mean value of the summary statistic (if we are testing CSR, the theoretical value is supplied by fun; otherwise we perform a separate set of nsim2 simulations, compute the average of all these simulated values, and take this average as an estimate of the theoretical mean value). Then, for each simulation, we compare the simulated curve to the theoretical curve, and compute the maximum absolute difference between them (over the interval of r values specified by ginterval). This gives a deviation value d[i] for each of the nsim simulations. Finally we take the m-th largest of the deviation values, where m=nrank, and call this dcrit. Then the simultaneous envelopes are of the form lo = expected - dcrit and hi = expected + dcrit where expected is either the theoretical mean value theo (if we are testing CSR) or the estimated theoretical value mmean (if we are testing another model). The simultaneous critical envelopes have constant width 2 * dcrit.

## "The simultaneous critical envelopes allow us to perform a different Monte Carlo test (Ripley, 1981). The test rejects the null hypothesis if the graph of the observed function lies outside the envelope at any value of r. This test has exact significance level alpha = nrank/(1 + nsim)."

## So for exact significance = 0.05, with nsim = B = 199, I need nrank = 10
## Or, with nsim = B = 499, I need nrank = 25

alpha <- 0.05 ### my desired global significance level
rank <- alpha * (1 + B) ### B was defined way toward the beginning of this file as the number of simulations to use

yKlimits <- c(-200, 200)
pdf(file = "SimulationKm12.pdf")
plot(envelope(m12, Kinhom, nsim = B, nrank = rank,  global = TRUE, ginterval = c(0,3), expression = project2segment(simulate(m12, nsim = 1, drop = TRUE), srems.roads.psp)$Xproj), xlim = c(0,3), main = "inhomogeneous K function", legend = FALSE, lwd = 2, ylim = yKlimits)
legend(legend=c("observed from cases", "theoretical from model", "95% simulation envelope"), lty = c(1, 2, 8), lwd = c(3,3,8), col = c("black", "red", "grey"), "topleft")
dev.off()
## ## and based on Ginhom
## not run
## pdf(file = "SimulationGm12.pdf")
## plot(envelope(m12, Ginhom, nsim = B, nrank = rank, global = TRUE, ginterval = c(0,3), expression = project2segment(simulate(m12, nsim = 1, drop = TRUE), srems.roads.psp)$Xproj), xlim = c(0,3), main = "inhomogeneous G function", legend = FALSE, lwd = 2, ylim = c(-0.1, 0.1))
## legend(legend=c("observed cases", "theoretical from model", "95% simulation envelope"), lty = c(1, 2, 8), lwd = c(3,3,8), col = c("black", "red", "grey"), "topleft")
## dev.off()

## and for comparison, from the null model with the offset as the only "predictor"
pdf(file = "SimulationKm0.pdf")
plot(envelope(m0, Kinhom, nsim = B, nrank = rank, global = TRUE, ginterval = c(0,3), expression = project2segment(simulate(m0, nsim = 1, drop = TRUE), srems.roads.psp)$Xproj), xlim = c(0,3), main = "inhomogeneous K function", legend = FALSE, lwd = 2, ylim = yKlimits)
legend(legend=c("observed from cases", "theoretical from model", "95% simulation envelope"), lty = c(1, 2, 8), lwd = c(3,3,8), col = c("black", "red", "grey"), "topleft")
dev.off()
## ## and based on Ginhom
## not run
## pdf(file = "SimulationGm0.pdf")
## plot(envelope(m0, Ginhom, nsim = B, nrank = 5, global = TRUE, ginterval = c(0,3), expression = project2segment(simulate(m0, nsim = 1, drop = TRUE), srems.roads.psp)$Xproj), xlim = c(0,3), main = "inhomogeneous G function", legend = FALSE, lwd = 2, ylim = c(-0.1, 0.1))
## legend(legend=c("observed cases", "theoretical from model", "95% simulation envelope"), lty = c(1, 2, 8), lwd = c(3,3,8), col = c("black", "red", "grey"), "topleft")
## dev.off()


###################################################
### code chunk number 22: rhohatMMdist
###################################################
### not run--probably too deep to get into for this audience
## ## is functional form for MMdist really linear?
## plot(parres(m12, "MMdist"), xlim = c(0,3), lwd = c(3,1,1,2), col = c("blue", "grey", "grey", "red"))
## ## does MMdist predict well across all its values?
pdf(file = "RhohatMMdist.pdf")
plot(rhohat(m12, MMdist), xlim = c(0,5), legend = FALSE, main = "underprediction near minimarts", xlab = "distance in km from nearest minimart")
abline(h = 1, col = "blue")
text(2, 5, labels = expression(lambda(u) == rho(MMdist) * lambda[0](u)), cex = 1.5)
text(2, 4, labels = expression(rho(MMdist) == lambda(u) / lambda[0](u)), cex = 1.5)
arrows(0.25, 0, 0.25, 1.6, col = "purple", lwd = 6)
dev.off()
## the following is not very helpful
## plot(rhohat(m12, popDens.f))
## abline(h = 1, col = "blue")


###################################################
### code chunk number 23: displaysingletermdroptablem12
###################################################
latex(singleTermDrop1table.m5, file = "",  rowname = c("(Intercept)", "Population density", "Poverty rate", "Owner-occupancy", "Percent age 20-44", "Distance to minimart"), caption = "Preliminary multi-predictor model. Df = degrees of freedom for a likelihood ratio test. AIC = Akaki's Information Criteria if the predictor was removed from the model. LRT = likelihood ratio test statistic. Pr(>Chi) = p-value for the likelihood ratio test.", digits = 2, rowlabel = "", booktabs = TRUE, label = "prelimmodel")


###################################################
### code chunk number 24: displaydeltabetahattable
###################################################
latex(percent.delta.betahat, file = "",  rowname = c("(Intercept)", "Population density", "Poverty rate", "Owner-occupancy", "Percent age 20-44", "Distance to minimart"), label = "deltabetahat", caption = "Percent change in remaining model coefficients upon deletions. Blank cell(s) in each column indicate the predictor that was removed. Column m11 indicates the effect of removing all non-significant predictors from the preliminary model simultaneously. The only removal that leaves the remaining coefficients little-changed is that of poverty rate, shown in column m7", rowlabel = "", digits = 2, booktabs = TRUE)


###################################################
### code chunk number 25: displaymodeltable
###################################################
latex(model.table, file = "", label = "finalmodel", rowlabel = "", rowname = c("(Intercept)", "Population density", "Owner-occupancy", "Percent age 20-44", "Distance to minimart"), cgroup = c("", latexTranslate("95%  CI")), n.cgroup = c(1,2), colheads = c("Odds ratio", "LB", "UB"), booktabs = TRUE, caption = "Final model. CI = confidence interval. LB = lower bound. UB = upper bound.")
